# Leveraging transcript quantification for fast computation of alternative splicing profiles

**DOI:** 10.1101/008763

**Authors:** Gael P. Alamancos, Amadís Pagès, Juan L. Trincado, Nicolás Bellora, Eduardo Eyras

**Affiliations:** Universitat Pompeu Fabra, E08003, Barcelona, Spain; Centre for Genomic Regulation, E08003, Barcelona, Spain; INIBIOMA, CONICET-UNComahue, Bariloche, Río Negro, Argentina; Catalan Institution for Research and Advanced Studies, E08010 Barcelona, Spain

**Keywords:** RNA-Seq, splicing, splicing event

## Abstract

Alternative splicing plays an essential role in many cellular processes and bears major relevance in the understanding of multiple diseases, including cancer. High-throughput RNA sequencing allows genome-wide analyses of splicing across multiple conditions. However, the increasing number of available datasets represents a major challenge in terms of computation time and storage requirements. We describe SUPPA, a computational tool to calculate relative inclusion values of alternative splicing events, exploiting fast transcript quantification. SUPPA accuracy is comparable and sometimes superior to standard methods using simulated as well as real RNA sequencing data compared to experimentally validated events. We assess the variability in terms of the choice of annotation and provide evidence that using complete transcripts rather than more transcripts per gene provides better estimates. Moreover, SUPPA coupled with *de novo* transcript reconstruction methods does not achieve accuracies as high as using quantification of known transcripts, but remains comparable to existing methods. Finally, we show that SUPPA is more than 1000 times faster than standard methods. Coupled with fast transcript quantification, SUPPA provides inclusion values at a much higher speed than existing methods without compromising accuracy, thereby facilitating the systematic splicing analysis of large datasets with limited computational resources. The software is implemented in Python 2.7 and is available under the MIT license at https://bitbucket.org/regulatorygenomicsupf/suppa.

**Supplementary Information:** available at https://bitbucket.org/regulatorygenomicsupf/suppa/downloads/Supplementary_Data.zip

## Introduction

Alternative splicing plays an important role in many cellular processes and bears major relevance in the understanding of multiple diseases, including cancer (David & Manley 2010, Ward & Cooper 2010). Numerous genome wide surveys have facilitated the description of the alternative splicing patterns under multiple cellular conditions and disease states. These studies are generally based on the measurement of local variations in the patterns of splicing, encoded as events, and have carried out using microarrays (Thorsen et al. 2008, Lapuk et al. 2010, Misquitta-Ali et al. 2011), RT-PCR platforms (Klinck et al. 2008, Simpson et al. 2008), or RNA sequencing (Pan et al. 2008, Wang et al. 2008). The description of alternative splicing in terms of events facilitates their experimental validation using PCR methods and the characterization of regulatory mechanisms using sequence analysis and biochemical approaches (Bechara et al. 2013, Raj et al. 2014); and they provide a valuable description for predictive and therapeutic strategies (Xiong et al. 2014, Hua et al. 2015). Events are generally defined as local variations of the exon-intron structure that can take two possible configurations, and are characterized by an inclusion level, also termed PSI or Ψ, which measures the fraction of mRNAs expressed from the gene that contain an specific form of the event (Venables et al. 2008, Wang et al. 2008). In terms of sequencing reads, Ψ is usually defined as the ratio of the density of inclusion reads to the sum of the densities of inclusion and exclusion reads (Wang et al. 2008, Shen et al. 2012). Initial methods to estimate Ψ values were based on reads from junction, exons or both (Pan et al. 2008, Wang et al. 2008, Sultan et al. 2008). Later methods were developed that take into account the uncertainty of quantification from single experiments (Katz et al. 2010), the comparison of two conditions (Katz et al. 2010, Griffith et al. 2010, Shen et al. 2012, Wu et al. 2011, Shi et al. 2013), as well as multiple replicates per condition (Shen et al. 2012, Brooks et al. 2011, Singh et al. 2011, Hu et al. 2013) and paired-replicates (Shen et al. 2014).

Current tools to process RNA sequencing data to study alternative splicing events can take more than a day to analyze a single sample and often require excessive storage, so they are not competitive to be applied systematically to large data sets, unless access to large computational resources is granted. In particular, methods for estimating Ψ values generally involve the mapping of reads to the genome or to a library of known exon-exon junctions, both of which require considerable time and storage. Additionally, accuracy is often achieved at the cost of computing time. All this represents a major obstacle for the analysis of large datasets, and in particular, for the re-analysis of public data and updates with new annotations or assembly versions. More importantly, these analyses remain unfeasible at small labs with limited computational resources. On the other hand, recent developments in the quantification of known transcripts have shown that considerable accuracy can be achieved at high speed (Li et al. 2011, Roberts et al. 2013, Patro et al. 2014, Zhang et al. 2014). This raises the question of whether fast transcript abundance computation could be used to obtain accurate estimates of Ψ values for local alternative splicing events genome wide.

In this article we describe SUPPA, a computational tool to leverage fast transcript quantification for rapid estimation of Ψ values directly from the abundances of the transcripts defining each event. Using simulated data we show that Ψ values estimated by SUPPA, coupled to Sailfish or RSEM transcript quantification, are closer to the ground-truth than two standard methods, MATS and MISO. Additionally, using an experimentally validated set and matched RNA-Seq data we show that SUPPA achieves slightly superior or comparable accuracy compared with MATS and MISO. We further assess the variability in terms of the choice of annotation and provide evidence that using complete transcripts rather than more transcripts per gene in the annotation provide better estimates. Moreover, we show that SUPPA coupled with *de novo* transcript reconstruction methods does not achieve accuracies as high as using the quantification of known transcripts, but remains comparable to existing methods. Finally, speed benchmarking provides evidence that SUPPA can obtain Ψ values at a much higher speed than existing methods. We argue that coupled to a fast transcript quantification method, SUPPA provides a fast and accurate approach to systematic splicing analysis. SUPPA facilitates the accurate splicing analysis of large datasets, making possible for labs with limited computational resources to exploit data from large genomics projects and contribute to the understanding of the role of alternative splicing in cell biology and disease.

## Results

### SUPPA

SUPPA provides an effective and easy-to-use software to calculate the inclusion levels (Ψ) of alternative splicing events exploiting transcript quantification (Figure 1A). An alternative splicing event is a local summary representation of the exon-intron structure from the transcripts that cover a given genic region, and is generally represented as a binary form, although more complex variations may happen. Accordingly, an event can be characterized in terms of the sets of transcripts that describe either form of the event, which can be denoted as *F*_1_ and *F*_2_. For instance, for an exon-skipping event, *F*_1_ represents the transcripts that include the exon, whereas *F*_2_ represents the transcripts that skip the exon. The inclusion value (Ψ) of an event is defined as the ratio of the abundance of transcripts that include one form of the event, *F*_1_, over the abundance of the transcripts that contain either form of the event, *F*_1_∪*F2* (Venables et al. 2008, Wang et al. 2008, Katz et al. 2010, Shen et al. 2012). Given the abundances for all transcripts isoforms, assumed without loss of generality to be given in transcript per million units (TPM) (Li et al. 2010), which we denote as *TPM_k_*, SUPPA then calculates Ψ for an event as follows:

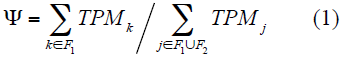

**Figure 1.**
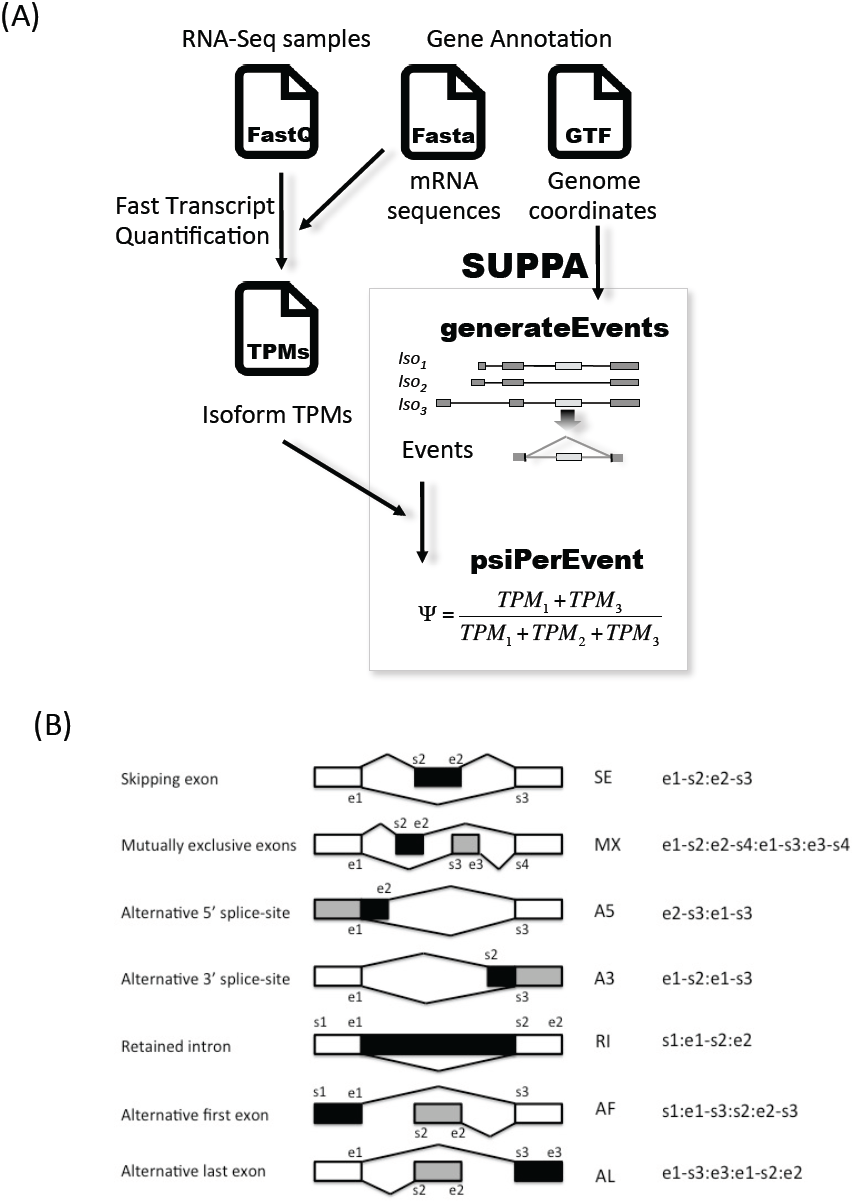
SUPPA pipeline. **(A)** SUPPA calculates possible alternative splicing events with the operation *generateEvents* from an annotation, which can be obtained from a database or built from RNA-Seq data using a transcript reconstruction method. For each event, the transcripts contributing to either form of the event are stored and the calculation of the Ψ value per sample for each event is performed using the transcript abundances per sample (TPMs) (Methods). From one or more transcript quantification files, SUPPA calculates for each event the Ψ value per sample with the operation *psiPerEvent*. SUPPA can use transcript quantification values obtained from any method. **(B)** Events generated from the annotation are given an unique identifier that includes a code for the event type (SE, MX, A5, A3, RI, AF, AL) and a set of start (s) and end (e) coordinates that define the event (shown in the figure) (Methods). In the figure, the form of the alternative splicing event that includes the region in black is the one for which the relative inclusion level (Ψ) is given: for SE, the PSI indicates the inclusion of the middle exon; for A5/A3, the form that minimizes the intron length; for MX, the form that contains the alternative exon with the smallest start coordinate (the left-most exon) regardless of strand; for RI, the form that retains the intron; and for AF/AL, the form that maximizes the intron length. The gray area denotes the alternative form of the event.

SUPPA provides the identifiers for the transcripts that describe either form of the event, which in combination with the transcript quantification is used to obtain the Ψ values using formula (1) (Figure 1A). SUPPA is agnostic of the actual methodology for quantifying transcripts and can read the quantification from multiple experiments from a single input. SUPPA generates different alternative splicing events types from an input annotation file in GTF format: exon skipping (SE), alternative 5’ and 3’ splice-sites (A5/A3), mutually exclusive exons (MX), intron retention (RI), and alternative first and last exons (AF/AL) (Figure 1B). The Ψ value for an event is calculated with respect to one of the two forms of the event (Figure 1B). Further details and options of the software are given at https://bitbucket.org/regulatorygenomicsupf/suppa/.

### Accuracy analysis with simulated data

Transcript abundances and corresponding paired-end reads were simulated using FluxSimulator (Griebel et al. 2012) with the RefSeq annotation as reference (Methods) (Supplementary Table 2). The reference set for accuracy analysis was built using events in genes with only two alternative transcripts in the RefSeq annotation that did not overlap any other events. In these cases, the Ψ of the event is identical to the relative abundance of one of the two transcripts. The ground-truth Ψ values were then defined to be the relative abundances of the transcripts isoforms in these genes, where the transcript abundances were taken to be the simulated abundances. Simulated RNA-Seq reads were mapped to the genome and used to calculate Ψ_MISO_ and Ψ_MATS_ values with MISO (Katz et al. 2010) and MATS (Shen et al. 2011), respectively (Methods). The same simulated reads were also used to quantify transcript abundances with Sailfish (Patro et al. 2014) and RSEM (Li et al. 2011), and Ψ_Sailfish_ and Ψ_RSEM_ values were then calculated with SUPPA (Methods). Only genes with total transcripts per million (TPM) abundance, calculated as the sum of the TPM of its transcripts, greater than 1 were considered. This resulted in a set of 144 events (Supplementary Data 1). Comparing the four sets of estimated Ψ values with the ground-truth, the Ψ_Sailfish_ and Ψ_RSEM_ values calculated with SUPPA show the highest correlations (Table 1) (Figure 2A). Moreover, calculating how different the estimated Ψ values are from the ground-truth, SUPPA Ψ values (Ψ_Sailfish_ and Ψ_RSEM_) show the closest behaviour, followed by MISO and MATS, which behave similarly (Figure 2B).

**Figure 2.**
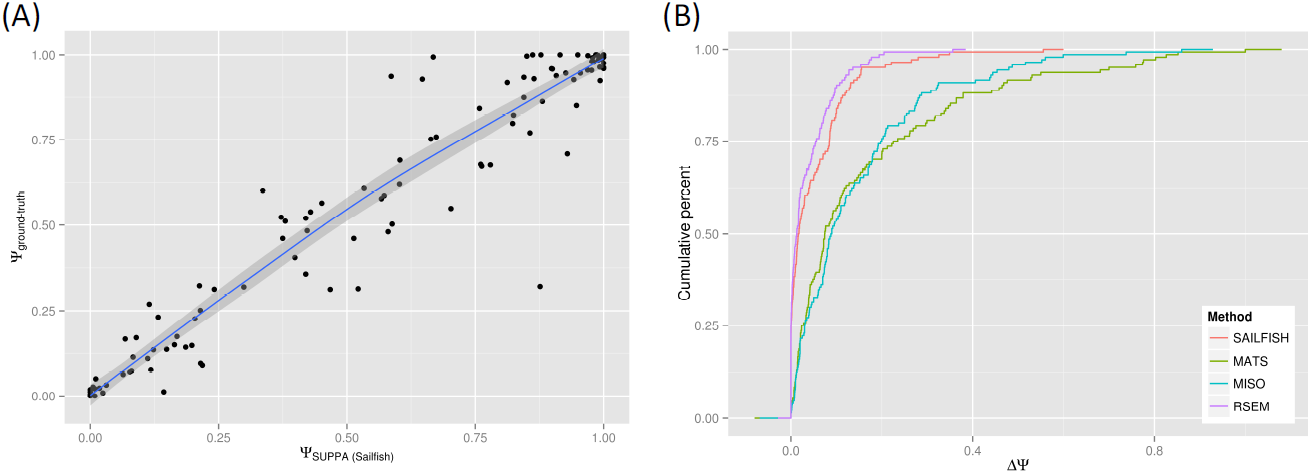
Benchmarking with simulated data. **(A)** Correlation of the ground-truth Ψ values (Methods) with those estimated with Sailfish+SUPPA using simulated data. The blue line and gray boundaries are the fitted curves with the LOESS regression method. **(B)** Cumulative distribution of the absolute difference between the ground-truth Ψ values and the ones estimated with Sailfish+SUPPA (SAILFISH), RSEM+SUPPA (RSEM), MISO and MATS. The lines describe the proportion of all events tested (Cumulative percent, y-axis) that are predicted at a given maximum absolute difference from the ground-truth value (ΔΨ, x-axis).

**Table 1.** First row: Correlation values (Spearman and Pearson) between the estimated and ground-truth Ψ values using simulated data. The comparison involves 144 events (Supplementary Data 1). Second and third rows: correlation values (Spearman and Pearson R) between the estimated Ψ values from ESRP1-overexpressed (ESRP) and empty-vector (EV) RNA-Seq datasets and the RT-PCR validation for the same samples (Shen et al. 2012). This comparison involves the 60 events that were in the RefSeq annotation and had a Ψ value from every method (Supplementary Data 2).

| *Annotation* | *Dataset* | **Sailfish+SUPPA** |  | **RSEM+SUPPA** |  | **MATS** |  | **MISO** |  |
| --- | --- | --- | --- | --- | --- | --- | --- | --- | --- |
|  |  | *Pearson* | *Spearman* | *Pearson* | *Spearman* | *Pearson* | *Spearman* | *Pearson* | *Spearman* |
| ***RefSeq*** | ***Synthetic*** | 0,971 | 0,959 | 0,987 | 0,978 | 0,833 | 0,819 | 0,862 | 0,815 |
|  | ***ESRP1*** | 0,778 | 0,769 | 0,795 | 0,779 | 0,763 | 0,753 | 0,701 | 0,715 |
|  | ***EV*** | 0,767 | 0,766 | 0,808 | 0,823 | 0,805 | 0,815 | 0,765 | 0,782 |
| ***Ensembl*** | ***ESRP1*** | 0,633 | 0,627 | 0,682 | 0,673 | 0,723 | 0,715 | 0,708 | 0,691 |
|  | ***EV*** | 0,608 | 0,613 | 0,664 | 0,668 | 0,794 | 0,790 | 0,747 | 0,774 |

### Accuracy analysis with experimentally validated events

To further validate the calculation of Ψ values with SUPPA, we used a set of 163 alternative splicing events validated by RT-PCR in MDA-MB-231 cells under two conditions: with overexpression of the splicing factor ESRP1 (ESRP1) and with an empty vector (EV) (Shen et al. 2012). We used the RNA-Seq data obtained from the same samples (Shen et al. 2012) to predict the Ψ values as before. From both RNA-Seq datasets we quantified the RefSeq transcripts using Sailfish and RSEM, and calculated the SUPPA Ψ_Sailfish_ and Ψ_RSEM_ values. RNA-Seq reads were mapped to the genome to run MISO and MATS to obtain the corresponding Ψ values (Methods). From the 163 validated events, we finally compared those 60 that were present in the RefSeq annotation and for which we had Ψ values for all methods (Supplementary Data 2). Sailfish+SUPPA and RSEM+SUPPA show an overall slightly better correlation than the other methods for the ESRP1 sample, whereas RSEM+SUPPA and MATS show the best correlations for the EV sample (Table 1) (Figure 3A). Although RSEM+SUPPA shows the highest correlations in all cases, Sailfish+SUPPA correlations are comparable to the rest. Calculating the absolute difference between the estimated and the experimental Ψ values for each event, we observe that SUPPA, either combined with Sailfish or RSEM, is more accurate than MISO and MATS (Figure 3B). Performing the same analysis using the Ensembl annotation, comparing a total of 91 events common to all approaches, we observe a general decrease of accuracy in all methods (Table 1) (Supplementary Figure 1).

**Figure 3.**
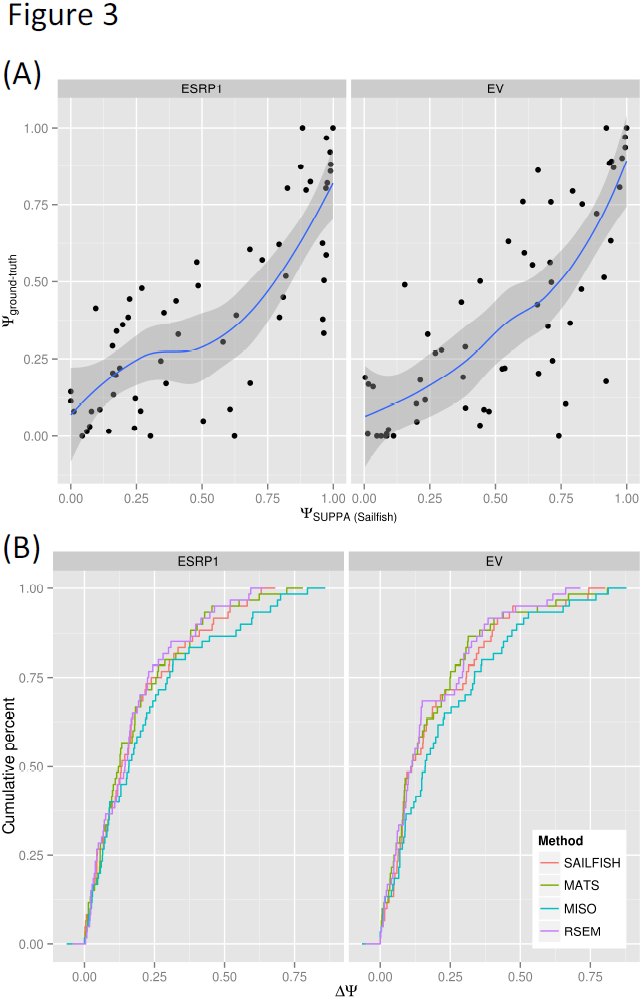
Benchmarking using experimentally validated events. **(A)** Correlation of the experimental Ψ values with those estimated with Sailfish+SUPPA in MDA-MB-231 cells with (ESRP1, left panel) and without (EV, right panel) ESRP1 overexpression. Experimental Ψ values were obtained using RT-PCR (Shen et al. 2012) and estimated Ψ were obtained from RNA-Seq data from the same samples (Shen et al. 2012). The blue line and gray boundaries are the fitted curves with the LOESS regression method. **(B)** Cumulative distribution of the absolute difference between the same experimental Ψ values and the ones estimated with Sailfish+SUPPA (SAILFISH), RSEM+SUPPA (RSEM), MISO and MATS from RNA-Seq data from the same samples (Shen et al. 2012). The lines describe the proportion of all events (Cumulative percent, y-axis) that are calculated at a given maximum absolute difference from the RT-PCR value (ΔΨ, x-axis).

### Variability associated to replicates and annotation choice

To study how the choice of annotation may impact the accuracy of Ψ estimation, we obtained RNA from two biological replicates for the cytosolic fractions of MCF7 and MCF10 cells and performed sequencing using standard protocols. Correlation between replicates of the SUPPA Ψ values, using quantification with Sailfish on the RefSeq annotation, is high in all comparisons (Person R ~0.86-0.89) (Supplementary Figure 2). Furthermore, restricting this analysis to genes with TPM>1, calculated as the sum of the TPMs from all transcripts in each gene, the correlation between replicates increases (Pearson R~0.95-0.97) (Supplementary Figure 2). We then compared the results obtained using SUPPA with the quantifications on the RefSeq and Ensembl transcripts. SUPPA Ψ values were calculated using both replicas of the cytosolic MCF7 RNA-Seq data (similar results were observed for MCF10, data not shown). The comparisons were performed using the 9301 (MCF7, replica 1) and 9287 (MCF7, replica 2) events that were found in both annotations, and not overlapping with other events from the same replica. We observe variability in the estimation of Ψ between annotations that does not depend on the difference in the number of transcripts used for Ψ calculation (Figure 4A). Similarly, this variability is also independent of the difference in the number of transcripts annotated in the gene in which the event is contained (Supplementary Figure 3). Moreover, the disparity in Ψ estimates is also independent of the mean expression of the gene in which the event is contained (Figure 4B). On the other hand, the dispersion of Ψ estimates comparing replicas and using the same annotation decreases with the mean expression of the gene (Figure 4C), which at low expression it is comparable to the dispersion for Ψ estimates as a result of differences in annotation (Figures 4A and 4B).

**Figure 4.**
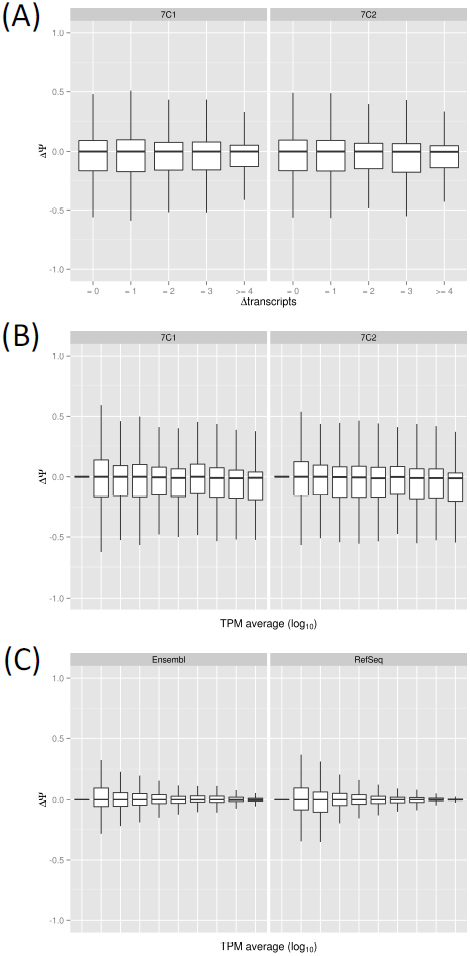
Annotation dependencies. Boxplots of the difference of Ψ values estimated by SUPPA for Ensembl and RefSeq annotations from Sailfish quantification (y axis) as a function of **(A)** the difference in the number of transcripts defining each event in Ensembl and RefSeq or as a function of **(B)** the mean expression of the gene in which the event is contained. The x-axis in (B) is grouped into 10 quantiles according in the log10(TPM) scale. The variability is represented for both replicates (7C1 and 7C2) of the cytosolic RNA-Seq data from MCF7 cells. **(C)** Boxplots of the distribution of Ψ differences between replicates for the estimates from the Ensembl (left panel) and RefSeq (right panel) annotations as a function of the mean expression genes (x-axis), grouped into 10 quantiles in the log_10_(TPM) scale, using genes with TPM>0. Mean expression is calculated as the average of the log_10_(TPM) for the each gene in the two replicates for (C) or for the each gene in the two annotations in (B).

### Annotation-free estimation

The previous analyses suggest that incomplete annotations may lead to inaccurate transcript quantification, which will have in turn a negative impact on the Ψ estimates by SUPPA. Methods for *de novo* transcript reconstruction facilitate the discovery of new transcripts missing from the annotation and the completion of existing ones from RNA-Seq reads (Trapnell et al. 2010, Li et al. 2011b, Li et al. 2011c, Li et al. 2012, Mezlini et al. 2012, Behr et al. 2013, Tomescu et al. 2013, Rossell et al. 2014, Maretty et al. 2014). As these methods produce an annotation of transcripts and their corresponding abundances, their output can be used with SUPPA to calculate alternative splicing events and their Ψ values. They thus provide an opportunity to assess whether a *de novo* prediction of transcripts structures and subsequent quantification from RNA-Seq data may lead to more accurate Ψ values than using a fixed annotation. To test this, we run Cufflinks with the *de novo* options with RNA-Seq data from the ESRP1 and EV samples (Methods). Using the resulting annotation, we calculated all possible alternative splicing events and their contributing transcripts with SUPPA. Similarly, we calculate the Ψ with MATS and MISO using the same reads mapped to the genome, this time guided by the Cufflinks annotation. We then compared the Ψ values obtained for the events in common with the experimentally validated set (Shen et al. 2012): 82 for ESRP1 and 47 for EV (Supplementary Data 3). We observe that for all approaches the correlation of Ψ values decreases (Table 2). The Ψ_Cufflinks_ values obtained with SUPPA (Figure 5) (Table 2) are comparable to the values obtained using the Ensembl annotation (Table 1). Moreover, we recalculated the transcript quantification using Sailfish on the Cufflinks annotations, but found no improvement (Table 2).

**Figure 5.**
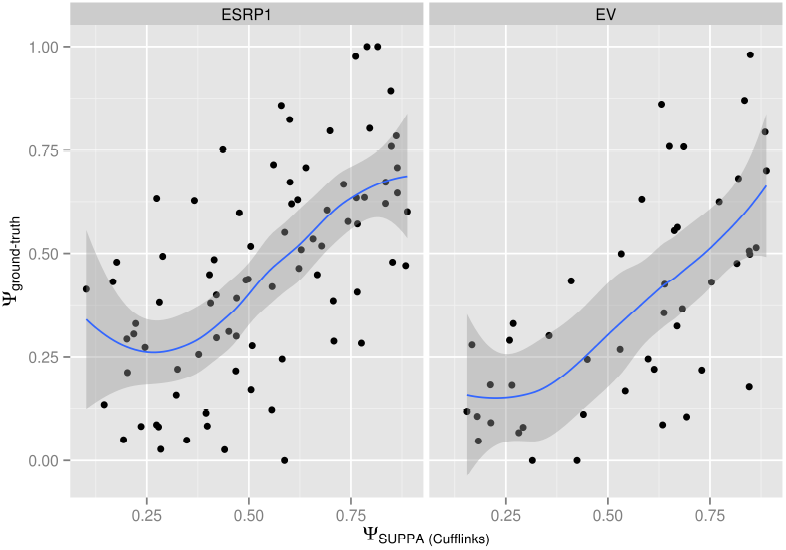
Annotation-free PSI estimation. Correlation of the experimental Ψ values with those estimated with Cufflinks *de novo* + SUPPA in MDA-MB-231 cells with (ESRP1, left panel) and without (EV, right panel) ESRP1 overexpression. Experimental Ψ values were obtained by RT-PCR (Shen et al. 2012) and estimated PSIs were obtained from RNA-Seq data in the same samples (Shen et al. 2012). The blue line and gray boundaries are the fitted curves using the LOESS regression method.

**Table 2.**
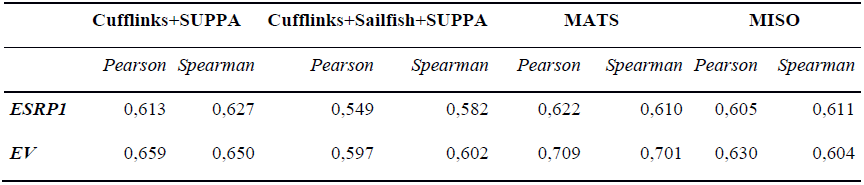
Correlation values (Spearman and Pearson R) between the estimated Ψ values from ESRP1-overexpressed (ESRP) and empty-vector (EV) RNA-Seq datasets and the RT-PCR validation for the same samples (Shen et al. 2012). This comparison involves 83 events in the ESRP1 sample and 47 in the EV sample, which can be found in Supplementary Data 3.

|  | Cufflinks+SUPPA |  | Cufflinks+Sailfish+SUPPA |  | MATS |  | MISO |  |
| --- | --- | --- | --- | --- | --- | --- | --- | --- |
|  | *Pearson* | *Spearman* | *Pearson* | *Spearman* | *Pearson* | *Spearman* | *Pearson* | *Spearman* |
| **ESRP1** | 0,613 | 0,627 | 0,549 | 0,582 | 0,622 | 0,610 | 0,605 | 0,611 |
| **EV** | 0,659 | 0,650 | 0,597 | 0,602 | 0,709 | 0,701 | 0,630 | 0,604 |

### Speed benchmarking

The time needed by each methodology to obtain the Ψ values from a FastQ file depends on multiple different steps. To make a comparative assessment of computation times we broke down the benchmarking into three different tasks, equivalent to the three necessary steps for the SUPPA analysis. The first step involves the calculation of alternative splicing events from an annotation file, which only needs to be carried once for a give annotation. To calculate 66577 alternative splicing events from the Ensembl 75 annotation (37494 genes, 135521 transcripts), SUPPA *generateEvents* took 20 minutes, whereas to calculate 16714 alternative splicing events from the RefSeq annotation (25937 genes, 48566 transcripts), it took 3 minutes.

The second step consists in the assignment of reads to transcripts and/or genomic positions. For the purpose of speed benchmarking of read-assignment to transcripts, although transcript abundance estimation includes extra computation steps, we considered the transcript quantification by Sailfish to be approximately equivalent to the read mapping to a reference genome. To perform this speed comparison we used the synthetic data (45 millions of paired-end reads) and both (ESRP1 and EV) RNA-Seq samples from the MDA-MB-231 cells pooled together (256 millions of single-end reads), and used STAR (Dobin et al. 2012) and TopHat as a comparison. Sailfish and STAR are the fastest to assign reads to their likely molecular sources, compared to TopHat and RSEM (Figure 6A).

**Figure 6.**
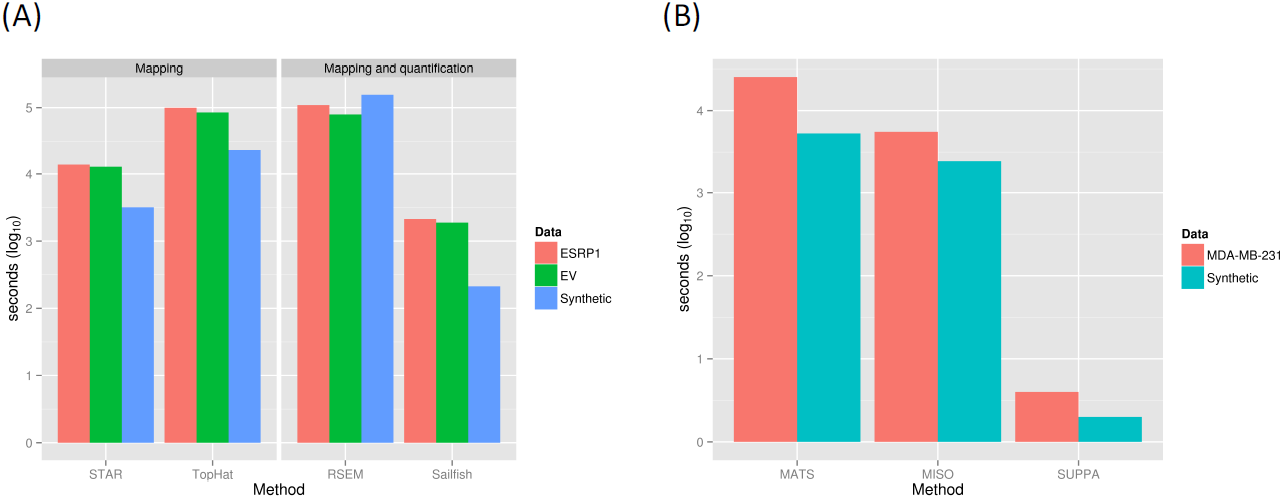
Speed benchmarking. **(A)**Time performance for read assignment/mapping to transcript/genome positions by RSEM, Sailfish, STAR and TopHat on the synthetic as well as the ESRP1 and EV RNA-Seq datasets separately (Methods). RSEM and Sailfish include the transcript quantification operation. **(B)** Time performance for the Ψ value calculation from the already mapped reads (MATS, MISO) or quantified transcripts (SUPPA). ESRP1 and EV samples were pooled for this benchmarking (MDA-MB-231). MATS time includes the calculation of the ΔΨ between samples, which we could not separate from the Ψ calculation (Shen et al. 2012). All tools were run in multi-threaded mode when possible. Time reported for all cases is the actual cumulative time the process used across all threads (Methods).

The third and final step is the Ψ calculation from either transcript quantification (SUPPA) or from the mapped reads (MISO and MATS). SUPPA *psiPerEvent* operation took less than a minute to produce an output size of 1Mb for 16714 events and was >1000 times faster than MISO and MATS on the same datasets (Figure 6B). In summary, the total time from the raw reads in FastQ format to the Ψ values for Sailfish + SUPPA against the RefSeq annotation-derived events took 214 (~ 3,5 mins) and 4022 (~ 1h) seconds for the synthetic and the MDA-MB-231 samples, respectively. We conclude that when used in conjunction with Sailfish, SUPPA is much faster than MISO and MATS, even if an ultra-fast aligner such as STAR (Dobin et al. 2012) is used for read mapping to the genome.

## Discussion

We have described SUPPA, a tool to calculate alternative splicing events from a given annotation and to estimate their Ψ values from the quantification of the transcripts that define the events. Using synthetic and experimental data, we have shown that SUPPA accuracy is generally comparable to and sometimes higher than other frequently used methods. Importantly, SUPPA can obtain Ψ values at a much higher speed without compromising accuracy. Moreover, SUPPA needs very little configuration, requires a small number of command lines for preprocessing and running and has no dependencies on Python libraries.

Although RNA-Seq data presents a number of systematic biases that need correction for accurate transcript quantification (Hansen et al. 2010, Li et al. 2010, Roberts et al. 2011), we did not observe differences in the accuracy of SUPPA when comparing corrected or uncorrected transcript quantification with Sailfish (data not shown). In fact, previous reports have already indicated that bias correction in RNA-Seq data does not influence the estimation of Ψ values (Shen et al. 2012, Zhao et al. 2013). On the other hand, we did observe that there is variability in the estimation of Ψ values associated to the choice of annotation. In the benchmarking using experimental data, using Ensembl annotation provides slightly worse accuracy than using RefSeq annotation, and this behaviour is consistent amongst all the tested methods. Interestingly, the observed variability between annotations does not depend on the difference in the number of transcripts per gene, on the number of transcripts used to describe the events, or on the expression of the gene in which the event is contained. On the other hand, the observed variability is comparable to the expected variability for lowly expressed genes between biological replicates. Such variability is in fact also frequently observed in transcript quantification methods (Patro et al. 2014, Maretty et al. 2014). It should be noted that RefSeq annotation includes less transcripts per gene than Ensembl, but these transcripts are mostly full-length mRNAs. In particular, RefSeq transcripts generally include complete untraslated regions, which generally hold a large contribution of the reads coming from a transcript, whereas a large proportion of Ensembl transcripts may be incomplete. These facts, together with our results, suggest that the completeness of the transcript structures, rather than the number of transcripts in genes, is determinant for an accurate estimate of transcript abundances, and consequently, for the correct estimate of event Ψ values with SUPPA.

Although SUPPA at the moment SUPPA generates the most common types of events, its model can be potentially expanded to more complex events, possibly involving more than two possible conformations. However, these complex events may not always be easy to test experimentally. On the other hand, the complexity may not always have to do with the number of possible conformations, but rather with a binary change that cannot be easily described in terms of just one or two exon boundary changes, as described recently for the gene QKI in lung adenocarcinomas (Sebestyen et al. 2015). We argue that a large proportion of the relevant splicing variation can be encapsulated with the binary events described by SUPPA and that more complex variations may be better described using transcript isoform changes (Sebestyen et al. 2015). Although SUPPA is limited to the splicing events available in the gene annotation, events can be expanded with novel transcript variants obtained by other means, like *de novo* transcript reconstruction and quantification methods. In this case we observed accuracies similar to the tests performed with the Ensembl annotations but lower than when using the RefSeq annotations. Moreover, performing quantification on the reconstructed transcripts using a different method does not improve the accuracy, indicating there is still a limitation on how well we can recover the right exon-intron structures *de novo* from RNA-Seq.

As transcript reconstruction and quantification methods improve in accuracy and methods for RNA sequencing increase their efficiency and reliability, our knowledge of the census of RNA molecules in cells will keep on progressing. Although single molecule sequencing methods may eventually lead to the abandonment of transcript reconstruction methods, they are still costly and error prone, and quantification still relies on short read sequencing. Transcript quantification methods will therefore continue to be an essential component in the description of the abundance of RNA molecules in cells. As fast reliable methods still depend on the annotation, future efforts may perhaps focus on improving transcript annotations under multiple conditions. In parallel to these advances, the local description of alternative splicing in terms of events will remain a valuable description RNA variability in genes in the context of studies of RNA regulation (Bechara et al. 2013, Raj et al. 2014) and of predictive and therapeutic strategies (Xiong et al. 2014, Hua et al. 2015).

In summary, when coupled to a fast transcript quantification method, SUPPA outperforms other methods in speed without compromising the accuracy. This is of special relevance when analyzing large amount of samples. Accordingly, SUPPA facilitates the systematic analyses of alternative splicing in the context of large-scale projects using limited computational resources. We conclude that SUPPA provides a method to leverage fast transcript quantification for efficient and accurate alternative splicing analysis for a large number of samples.

## Methods

### Alternative splicing events

The Ensembl annotation (Release 75) (Flicek et al. 2014) and the RefSeq annotation (NM_ and NR_ transcripts) (Pruitt et al. 2014) (assembly hg19) were downloaded in GTF format from the Ensembl FTP server and the UCSC genome table browser, respectively. All annotations on chromosomes other than autosomes or sex chromosomes were removed. In total, 37,494 genes and 135,521 transcripts were obtained for the Ensembl annotation, while 25,937 genes and 48,566 transcripts were obtained for the RefSeq annotation. We applied SUPPA to each annotation to obtain 16714 and 66577 events from RefSeq and Ensembl, respectively, including exon skipping (SE), alternative 5’ and 3’ splice-sites (A5/A3), mutually exclusive exons (MX), and intron retention (RI) events (Supplementary Table 1). Alternative first (AF) and last exons (AL) were not included in the analysis but can be also computed with SUPPA. Each event has a unique identifier that includes the gene symbol, the type of event, and the coordinates and strand that characterize the event:

~~~
*<gene_id>;<event_type>:<seqname>:<coordinates_of_the_event>:strand*
~~~

where *gene_id, seqname* and *strand* are obtained directly from the input annotation in GTF, *seqname* is the field 1 from the GTF file, generally the chromosome. The field *coordinates_of_the_event* is defined by start and end coordinates that define the *event_type* (SE, MX, A5, A3, RI, AF, AL).

### RNA sequencing data

A total of 45 million 2x50bp paired-end simulated reads were generated using FluxSimulator (Griebel et al. 2012) (parameter file described in Supplementary Table 2). RNA sequencing data from (Shen et al. 2012) was also used, corresponding to ESRP1-overexpression (ESRP1) and empty-vector (EV) experiments in MBA-MD-231 cells, available from the short read archive (SRA) under id SRX122589. Moreover, RNA sequencing was also performed in duplicate on cytosolic fractions of MCF7 and MCF10 cells using standard protocols (Supplementary Material), available at SRA under id SRP045592.

### Read mapping and PSI quantification

Read mapping to the genome was performed with the MATS pipeline (Shen et al. 2012), which uses TopHat (Trapnell et al. 2009) and an input annotation to map the reads. Reads mapping to *de novo* splice junctions were allowed, and those reads mapping to more than one genomic position were filtered out. For benchmarking, the same annotation used for transcript quantification was also used for read mapping to the genome in each of the comparisons (RefSeq, Ensemb or *de novo* Cufflinks). The mapping pipeline was run on simulated and real RNA-Seq reads. Mapped reads for each of the datasets were used with MATS, to obtain Ψ_MATS_ values for the different alternative splicing events (Supplementary Table 3). Similarly, mapped reads in SAM format were converted to BAM format and then sorted with samtools (Li et al. 2009) and analysed with MISO (Katz et al. 2010) to calculate the Ψ_MISO_ values for each of the datasets (Supplementary Table 4).

Sailfish (Patro et al. 2014) and RSEM (Li et al. 2011) were used to quantify all transcripts in the Ensembl and RefSeq annotations using the simulated and the real RNA-Seq datasets. The FASTA sequences of the transcripts corresponding to the same annotation as the GTF described earlier, were downloaded and used to generate the Sailfish index, selecting a *k*-mer size of 31 to minimize the number of reads assigned to multiple transcripts. Sailfish was then run using the FASTQ files for each read set and uncorrected and corrected (for sequence composition bias and transcript length) TPMs were calculated (Patro et al. 2014). RSEM was run as described previously (Li et al. 2011). The *psiPerEvent* operation of SUPPA was used to calculate the Ψ_Sailfish_ and Ψ_RSEM_ values from the transcript quantifications obtained by Sailfish and RSEM, respectively, for the alternative splicing events generated before, using the simulated and real datasets. The number of events for which SUPPA estimated a Ψ_Sailfish_ or Ψ_RSEM_ values are given in Supplementary Tables 5 and 6. For the purpose of benchmarking, the PSI values obtained from SUPPA (Ψ_Sailfish_ and Ψ_RSEM_), from MATS (Ψ_MATS_) and from MISO (Ψ_MISO_) for those events identified by all methods in each of the experiment, were compared with the simulated or the experimental values. Details of the commands used to run the different analyses are provided in Supplementary Tables 7-10. Supplementary data files with the alternative splicing events used in each on of the comparisons tested can be found at https://bitbucket.org/regulatorygenomicsupf/suppa/downloads/Supplementary_Data.zip

### Cufflinks analysis

The BAM files from 2 the MBA-MD-231 datasets were used to run Cufflinks (Trapnell et al. 2010) in order to generate and quantify transcriptome annotations *de novo*. The same read mapping as before was used. A total of 47211 transcripts were predicted and quantified for the ESRP1 dataset, whereas 37699 transcripts were predicted and quantified for the EV dataset. SUPPA *generateEvents* operation was then run on the GTF annotation generated by Cufflinks to calculate all the exon skipping events. This produced a total of 2566 and 2139 exon skipping events for the ESRP1 and EV datasets, respectively. Finally, SUPPA *psiPerEvent* operation was used to calculate the Ψ_Cufflinks_ values from the transcript quantification obtained by Cufflinks. For MISO and MATS, reads were mapped with the MATS pipeline using the Cufflinks annotation as input, and Ψ_MATS_ and Ψ_MISO_ were estimated as before. Additionally, Cufflinks reconstructed transcripts were used with SUPPA to quantify them from the same RNA-Seq data and to calculate Ψ values with SUPPA as before. The events common to all methods and coinciding with the experimentally validated ones were used for the benchmarking.

### Time benchmarking

All tools were run on the same node of an Oracle Grid Engine cluster, with 98Gb of RAM memory and 24 AMD Opteron (1.4 GHz) processors. All tools were run in multi-threaded mode when possible, but time reported is the actual cumulative time the process used across all CPUs.

## Acknowledgements

The authors wish to thank S. Mount, S. Janga, Y. Barash, M. Robinson for useful discussions and especially Y. Xing for useful discussions and for sharing sequencing and RT-PCR data. This work was supported by the Spanish Government [BIO2011-23920, CSD2009-00080], by the Sandra Ibarra Foundation for Cancer [FSI-2013] and partly by the Spanish National Institute of Bioinformatics (INB).

